# Coarse-tuning to improve a protein foundation model for a domain of proteins: the example of outer membrane beta barrels

**DOI:** 10.64898/2026.11.09.765412

**Authors:** Guy Yanai, Tomer Gan, Joanna S.G. Slusky, Margarita Osadchy, Rachel Kolodny

**Affiliations:** Department of Computer Science, University of Haifa, Haifa, Israel; Computational Biology Program and Department of Molecular Biosciences, University of Kansas, Kansas

**Keywords:** Protein foundation models training, Protein language models training, Transfer learning for proteins, Outer membrane beta barrels

## Abstract

Foundation models for proteins have shown great potential in addressing a wide range of challenges in computational molecular biology. State-of-the-art models are trained on large and diverse corpora of protein sequences, enabled by self-supervised training. The models are then used in transfer learning and the embeddings they calculate are used in supervised downstream tasks. We propose a novel focusing mechanism for foundation models, which we call ‘coarse-tuning’. In coarse-tuning, we further update the model, using a defined yet unlabeled, dataset to enhance its performance in a specific domain. We demonstrate this approach using Outer Membrane Beta Barrels (OMBBs) — proteins that are found in the distinct biophysical environment of the outer membrane. Our results show that coarse-tuned models outperform the original foundation model when applied to various transfer learning tasks. Additionally, we study the embeddings generated by the coarse-tuned models and compare them to those from the original foundation model, including an investigation of the learned attention matrices via contact map predictions. Our work suggests that a medium-sized dataset (fewer than 100,000 protein sequences) is sufficient to effectively specialize a foundation model to a specific domain, enhancing its utility for domain-specific applications.

## 1 Introduction

Contemporary machine learning techniques are transforming molecular biology and protein studies. To address newly defined challenges with deep learning tools, one needs not only to define the task, but also to collect datasets for model training and testing. Supervised training is easier as it requires less data, but supervised data must be labeled. For protein related tasks, foundation models that self-supervise their training using the databases of all protein sequences were shown to be very effective in that they capture general protein properties. These representations can then be used in downstream tasks.

Protein language models (PLM) are the prime example of foundation models that learn general protein properties. State-of-the-art [28] ESM exemplifies this, having learned from a dataset of 250 million proteins [26]. These models convert protein sequences to vectors or matrices that can be used by machine learning (ML) models [28, 4]. For example, the ML model can be a small network appended to the pre-trained PLM and trained in a supervised setting. If only the small network’s weights are adjusted, this is called transfer learning [23, 3]. If the weights of the PLM are also updated to optimize the loss on the supervised task, it is called fine-tuning (see Figure 1). Fine-tuning a large PLM with millions of parameters using a small dataset of thousands of examples can lead to overfitting. Separating representation learning from transfer learning can mitigate this issue.

**Fig 1.**
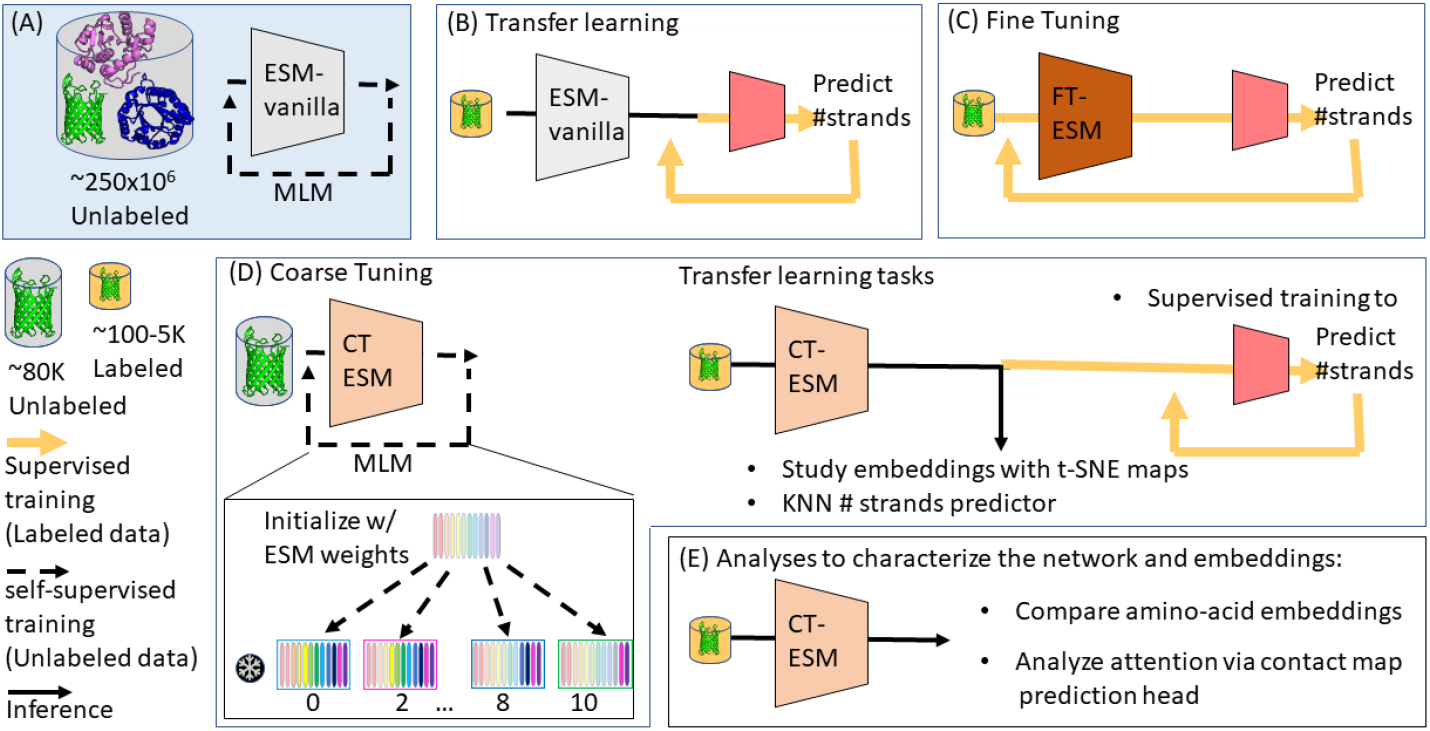
An overview of coarse-tuning a foundation model. (A) ESM-vanilla (unchanged-ESM2 model) was trained on MLM and a large and diverse set of proteins. Its embeddings can be used in a downstream task, which in our case is predicting the number of strands, in (B) transfer learning, so that only the small downstream network is trained (marked by color), or in (C) Fine-Tuning, so that the weights of both the small down-stream network and the foundation model are updated during the supervised training (marked by color change in both models). (D) We propose coarse-tuning the foundation model, using MLM and the set of ∼80,000 OMBB sequences. We consider different models that vary by the number of layers whose weights are frozen. We evaluate the coarse-tuned (CT) ESM foundation model by comparing it to the ESM-vanilla model on three transfer learning tasks, and (E) analyze directly the embeddings of the labeled set and the attention matrices via the contact map prediction head.

Masked language modeling (MLM) for training a foundation model only requires a dataset of sequences, as the network learns to predict randomly masked parts of the input. State-of-the-art PLMs, like ESM [26] and ProtTrans [10], are large transformer models trained with MLM. These models, whose training required extensive computation, are publicly available, and have proven effective for various downstream tasks. These include semantic similarities among proteins [30], predicting protein function [19], structure [17], and the effects of mutations [19, 2], classifying ion channels and ion transports [13]. For a recent review, see [28].

That PLMs effectively extract meaningful features from protein sequences is demonstrated by their utility in solving downstream tasks. Understanding what these features represent, remains a challenge. ESM’s vector values in the one-before-last layer of the model are the so-called ‘embeddings’, calculated for each residue in an input sequence as well as the <CLS> pre-pended and the <EOS> post-pended tokens. Two standard ML practices to convert the protein-length dependent embedding to fixed-size vectors are averaging over all residue embeddings, or taking (only) the embedding for the <CLS> token [5]. Both fixed-size representations depend on all residues: In the case of averaging this is straight-forward, and in the case of the <CLS> token via the models’ self-attention mechanism.

Insights into these embeddings can be gained by visualizing their relationships in lower dimensions, e.g., using t-SNE [18] as shown in the ESM study [26]. Vig et al. [29] and Rao et al. [24] also showed that the PLMs’ attention matrices are related to contacts in the protein structure. Because the attention matrices of ESM reveal information about the contacts in the input protein, the hugging-face [1] implementation allows their exploration.

We consider a setting where one studies a class of proteins, for which there is a relatively large non-labeled dataset (on the order of a few tens of thousands sample), but only a small labeled dataset (of a few hundred samples). As an example, we focus on outer membrane beta barrels (OMBB), an important class of membrane proteins [21, 9], that have these characteristics. OMBBs are found in Gram-negative bacteria, mitochondria, and chloroplasts and have diverse functions. Structurally, OMBBs are closed *β*-sheets, forming pores that traverse membranes [12] (see Supp. Figure 5 for examples). The sequence characteristics of OMBBs are distinct, both due to the unique biophysical constraints of the membrane environment imposed on these proteins, and the many evolutionary relationships among them [12, 11, 8]. We focus particularly on the most common OMBB types with 8 - 24 strands. Our non-labeled dataset includes ∼80,000 barrel sequences [20] and a few hundred proteins that are labeled by the number of strands in the barrel. For this setting, we propose using MLM to further train an ESM representation model – a process we call ‘coarse-tuning’, to enhance its performance on the downstream OMBB related tasks (see Figure 1). We evaluate the foundation model trained by coarse-tuning from several perspectives: the quality of the embeddings produced, the attention matrices learned, and the model’s effectiveness for an OMBB-related downstream task. Unlike the pre-training of ESM which relied on a huge set of 250 million sequences spanning the entire protein space, our coarse-tuning trains on a far smaller dataset of only ∼80,000 proteins. The improved performance of this refined model shows that medium-sized protein datasets can effectively tailor foundation models for specific domains.

## 2 Methods & Results

### 2.1 Training the coarse-tuned foundation model

We coarse-tuned a foundation model for OMBB related tasks and find that it performs better. The architecture and initial weights of our models are ESM2 with 35M parameters [26]. Our unlabeled dataset of ∼80,000 predicted OMBB sequences [20] is larger than typically used in classification tasks, albeit far smaller than the 250 million sequences used to train the initial ESM model. We selected this larger dataset over a smaller carefully-curated set with a higher accuracy of all proteins having the OMBB fold. The coarse-tuning training used the the Masked Language Modeling (MLM) task and the cross-entropy loss on 15% randomly chosen positions. Note that MLM is coherent even for the possibility of a few domains incorrectly labeled as OMBBs in our dataset, because our dataset only contains real domain sequences, so we do not mislead the model with false information. Moreover, training the model on a set of proteins containing primarily OMBBs and a small, and maybe even negligible, amount of non-OMBBs still serves the purpose of focusing ESM’s knowledge on OMBBs. We consider multiple models, which vary by the number of their initial frozen layers from the ESM models: 0, 2, 4, 6, 8, and 10 frozen layers (Fig. 1 (D)). We trained for 200 epochs on a random subset of 70% of the proteins in the dataset and monitored the cross-entropy loss on the remaining 30% (validation set).

The validation set has ∼24,600 OMBB proteins and was used (only) to monitor the training loss. Following ML best practices, the validation set is part, albeit distinct, of the training set, and is not included in any of the tests we perform on the resulting models. In all models the cross-entropy loss on the validation-set decreases and plateaus after 200 epochs (150000 steps). The performance of the different models trained with a different number of frozen layers in the MLM task was similar, in our following results we focus on the top-performing model in the chosen downstream tasks - the coarse-tuned model with no (i.e., 0) frozen layers. Suppl. Figure 2 shows the loss for the different models over training.

To study the impact of coarse-tuning the ESM model using OMBB data, we compare it to the unchanged-ESM2 model which we will refer to as ‘ESM-vanilla’. As comparing foundation models is challenging, we use multiple perspectives that compare their utility. In doing so, and although not our primary goal, comparing alternative perspectives provided us with insights into best-practice usage for foundation models. We noticed that calculating the embeddings by ESM-vanilla and the coarse-tuned model for padded inputs (used to resolve non-uniform input lengths in a batch), changes the embeddings significantly. Consequently, we avoided padded inputs by using a batch size of 1.

### 2.2 Probing the embeddings generated by coarse-tuned models

First, inspired by Figure 1 of the original ESM study [26], we compare the representations learned by the coarse-tuned and ESM-vanilla models by examining how they encode relationships between amino acid types. While the original study analyzed the learned output embedding weights, which provide a context-independent representation of each amino acid, we instead analyze contextualized residue embeddings. Specifically, for each model, we extract per-residue embeddings from a dataset of 102 outer membrane beta-barrel (OMBB) domains and average them across all occurrences of each amino acid type, obtaining a single representative embedding for each of the 20 amino acids. Unlike the learned output embeddings, these representations depend on both the model’s parameters and the dataset used to compute them, as each residue’s embedding is influenced by its surrounding sequence through the self-attention mechanism. Our approach therefore provides a dataset-conditioned view of how each model represents the different amino acid types in the context of OMBB proteins. To compare the relationships between amino acid types in the coarse-tuned and ESM-vanilla representation spaces, we visualize the resulting embeddings using t-SNE (Figure 2).

**Fig 2.**
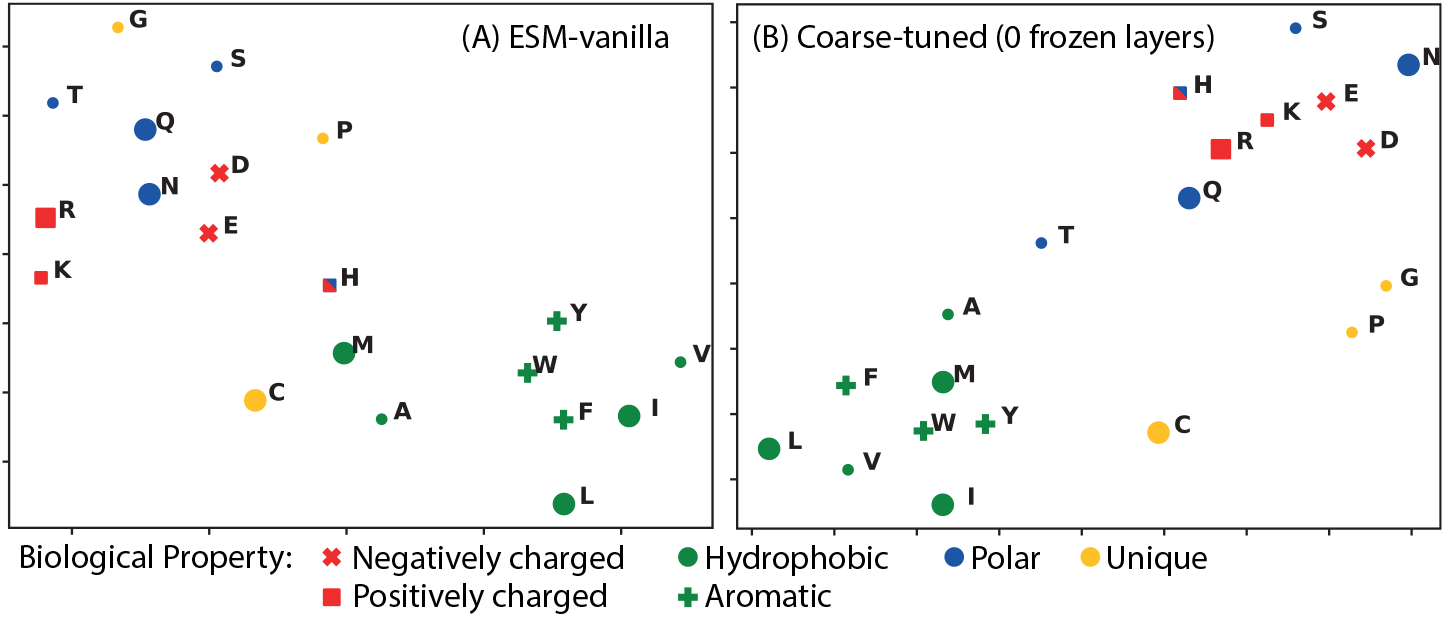
t-SNE embeddings of amino acids in ESM-vanilla and Coarse-tuned models on OMBB dataset

We find that coarse-tuning improves the biophysical relevance of the amino acid embeddings. This is especially true when considering hydrophobicity relationships among amino acids, potentially because of the opposite use of hydrophobicity for membrane barrels [6]. Specifically, compared to the t-SNE figure generated for the ESM-vanilla model (Figure 2A), in the coarse-tuned model’s figure (Figure 2A) we see that M is among the other hydrophobic residues, the position of A and T with respect to the hydrophobic and polar residues, Q and N are relatively further away from each other, compared to the positively charged R and K, and R and K are near the partially positively charged H and the negatively charged E and D. Thus we find that the relationships captured by the ESM-vanilla model for a set of OMBB proteins are subpar. In comparison, coarse-tuning the same model on OMBB sequences, i.e., by continuing the MLM training process on an OMBB dataset (which doesn’t include the domains used in this visualization), rescued the relationships among embeddings of different amino acids.

### 2.3 Evaluating the utility of the coarse-tuned model for transfer learning

We evaluate the utility of the coarse-tuned foundation model in transfer learning on the downstream task of predicting the number of strands in an OMBB (Figure 1 E). We compare two modes of transfer learning: the straightforward use of the embeddings ‘as-is’ and as inputs to a small network trained in a supervised setting. In both cases, we use a test set of proteins with the labels of their number of strands. For the case of supervised training a small network, we also use a labeled training set which is distinct from the above mentioned test set. We emphasize that the goal of correctly classifying the labeled test set is not used here in its own right but rather as a proxy to evaluate the utility of a foundation model.

Our test set includes 316 domains. 97 have experimentally solved structures, labeled by 8-24 strands (8 barrel classes altogether, see Supp. Figure 5) based on visual inspection of their structures [7]. 189 sequences whose structures were not solved experimentally, have structures predicted by AlphaFold, from which their labels were derived. 30 sequences are random non-barrel domains randomly sampled from ECOD (the ninth class).

We first compare directly the embeddings of the dataset labeled with the number of strands. Figure 3 shows the t-SNE visualizations of the mean domain embeddings for the test set proteins (calculated with a perplexity of 10 and 1000 iterations) of ESM-vanilla vs. the coarse-tuned model. The clustering is more pronounced in the coarse-tuned embeddings. Also, there are fewer cases of proteins with different classifications embedded near each other. To quantify this, we measure the accuracy of *k*-NN (*k* = 1, 3, 5) predictions using different distance measures (in high-dimensional embedding) between the vectors in a leave-one-out setting. Table 1 shows these measures, that are akin to listing the representation accuracy, and will be referred to as accuracy below. We find that in all cases, as measured by accuracy, the k-NN predictions based on the embeddings of the coarse-tuned model were measurably better than those of ESM-vanilla. Also, in all cases, the mean vector is a better representation of the full domain compared to the <CLS> vector.

**Table 1.** Results of 9-class classification of test set. The test set includes 97 OMBB domains with labels from their experimentally determined structures + 189 OMBB domains with labels from their AlphaFold predicted structures + 30 random nonbarrel domains (316 overall).

| **Model** | **1-NN Eu-clidean** | **3-NN Eu-clidean** | **5-NN Eu-clidean** | **1-NN Manhat-tan** | **3-NN Manhat-tan** | **5-NN Manhat-tan** | **1-NN Cosine** |
|---|---|---|---|---|---|---|---|
| **Embedding with <CLS> token** |  |  |  |  |  |  |  |
| ESM-vanilla | 88.92% | 88.29% | 88.29% | 88.29% | 88.29% | 87.97% | 88.92% |
| Coarse-tuned | 91.14% | 90.19% | 89.87% | 91.46% | 90.51% | 91.14% | 90.51% |
| **Embedding with mean vector** |  |  |  |  |  |  |  |
| ESM-vanilla | 91.14% | 91.46% | 90.82% | 92.09% | 92.72% | 92.41% | 91.14% |
| Coarse-tuned | 93.67% | 93.35% | 92.09% | 93.35% | 93.67% | 92.41% | 93.35% |

**Fig 3.**
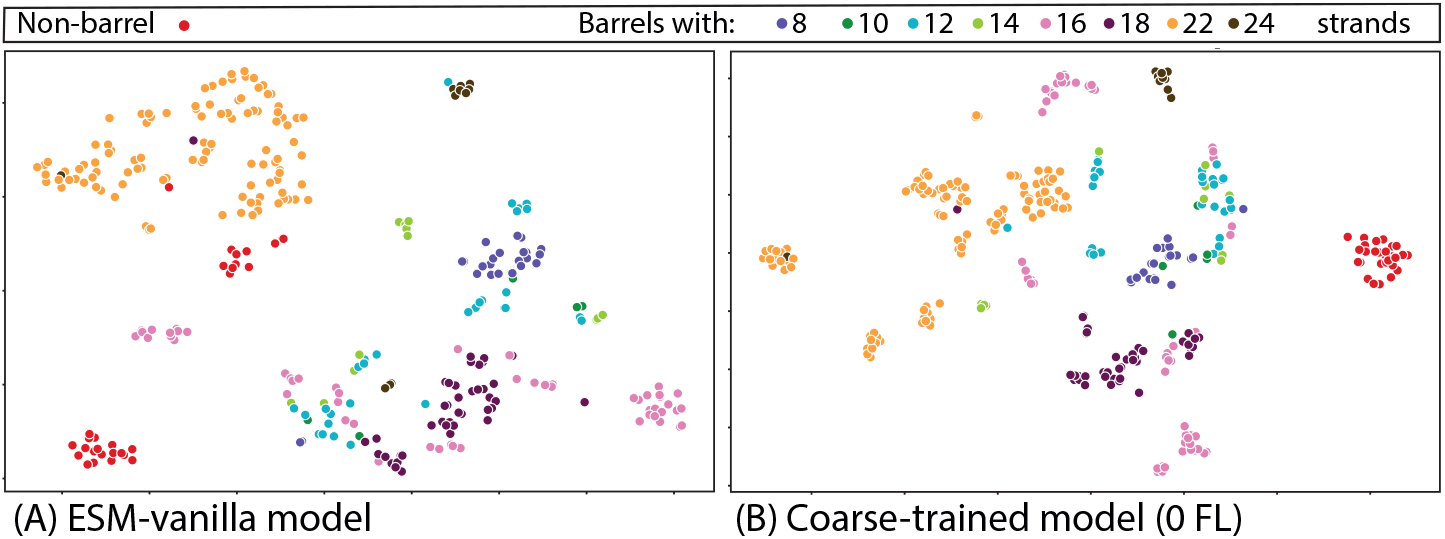
A t-SNE visualization of the sequence embeddings of the test set, consisting of 316 domains (286 OMBBs and 30 random, non-OMBB domains from ECOD): A) with ESM-vanilla model, B) with coarse-tuned model with 0 frozen layers

Following Radford et al.’s [22] approach, to further probe the embedding space of out coarse-tuned model, we trained small, fully connected 2-layer networks to predict the number of strands from the sequence embeddings. We intentionally used small networks that are by design weak classifiers. Hence, the success of the classifier network mainly depends on the quality of its input, which is the embedding of a protein, computed by the compared foundation models.

We trained the classifiers using an additional dataset of 8,739 OMBB proteins labeled by a computational pipeline [16] developed based on PolarBearal assignments of the AlphaFold predicted structures of the proteins’ sequences [7], split into training and validation sets with a ratio of 70:30. Supp. Figure 8 shows the validation loss during 40 epochs of supervised training. To evaluate the accuracies of the trained networks, we considered both our test set, and an expanded evaluation set which includes 291 designed (i.e., artificially generated) protein sequences. The designed sequences were generated by ESM-IF1 [15] (with default “temperature” parameter) given the beta-barrel structures of our labeled, experimentally-solved dataset. Three sequences were generated for each of the 97 barrels in said set. The labels for the designed sequences are the number of strands in the structure for which they were generated. We expect this expanded evaluation set to be harder to classify. Table 2 shows the accuracies of these trained networks on both sets, when using the mean and <CLS> embedding by the two foundation models.

**Table 2.** Comparison of Classification Performance for Different Models and Datasets.

| **Model** | **97 experimental structures** | **97 experimental + 189 AF-predicted structures** | **97 experimental + 189 AF-predicted + 30 non-barrel structures** | **291 OMBB designs by ESM-IF** |
|---|---|---|---|---|
| **Embedding with <CLS> token** |  |  |  |  |
| ESM-vanilla | 80.41% | 83.56% | 85.44% | 80.06% |
| Coarse-tuned | 91.75% | 90.55% | 90.82% | 87.62% |
| **Embedding with mean vector** |  |  |  |  |
| ESM-vanilla | 85.56% | 86.01% | 86.36% | 82.81% |
| Coarse-tuned | 92.78% | 89.86% | 89.86% | 90.37% |

We find that under all these tests, classifiers relying on the coarse-tuned model are more accurate than those relying on ESM-vanilla, and the accuracy gap can be up to 10%. This is true both for the labeled datasets of domains with known structure and the datasets with ESM-IF designed sequences. While not the focus of this study, we have a comparison of classifiers that differ in the input embedding: the <CLS> vector vs. the mean of the per-residue vectors. When comparing the embeddings directly, i.e., using the nearest neighbor classification, the mean vector is clearly a better representation. When we use a small network in a supervised setting (Table 2), and especially with the coarse-tuned foundation model, there are cases where the <CLS> embedding is better, and at any rate the difference in the achieved accuracy is smaller.

### 2.4 Using attention-based contact prediction as a proxy for foundation model comparison

Protein structure contact maps are two dimensional representations of the interactions between amino acids; here we consider a contact if their C-*α*s are closer than 8Å. Following [24], we consider protein structure contact prediction from the learned attention matrices in the coarse-tuned OMBB transformer model, and show that these are better predictors than the ones based on ESM-vanilla. Supplementary Figure 6 shows an example of the attention maps of the coarsetuned & ESM-vanilla models, compared to the true contact map for ECOD domain e1pt4A1 (an 8 stranded barrel). Contact maps of the beta-barrel fold have a distinctive pattern of perpendicular hatches along the diagonal, which are the contacts between neighboring strands. Also, there are the beta-seam contacts, the closing contacts between the first and last strands of the barrel (marked by a red box in the figure). The predicted contact maps of both models show some of this distinctive pattern, yet the coarse-tuned model better focuses its predictions on the ones in actual contact. Neither the attentions of the ESM-vanilla nor of the coarse-tuned model predict the beta-seam (and this is characteristic of our dataset).

**Fig 4.**
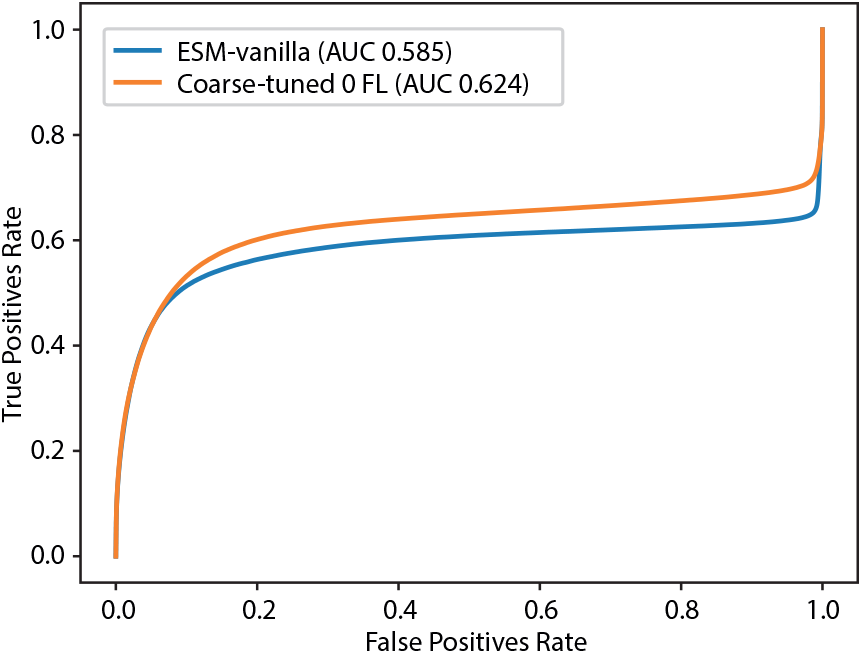
ROC curves for contact predictions on the dataset for ESM-vanilla and the coarse-tuned model. The ground truth is calculated from the C-*α* distances in the experimentally solved structures, and an 8Å threshold.

**Fig 5.**
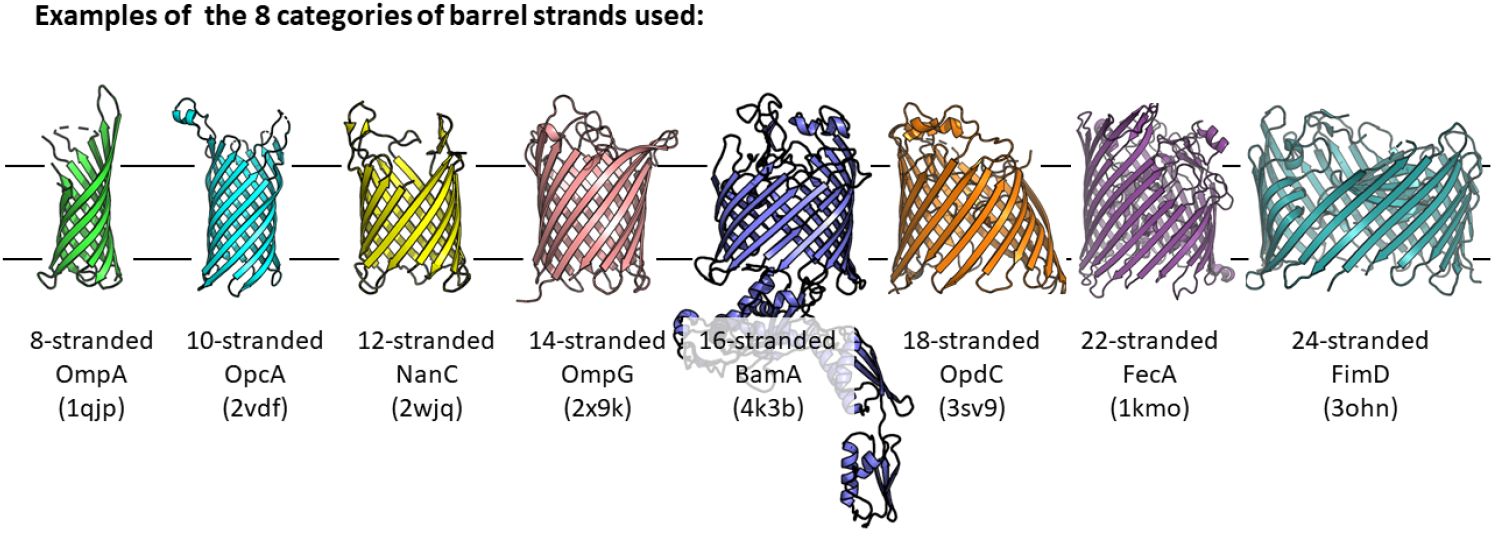
Examples of OMBB structures in each of the 8 classes with 8-24 strands. The structures are shown within their place in the membrane layer.

**Fig 6.**
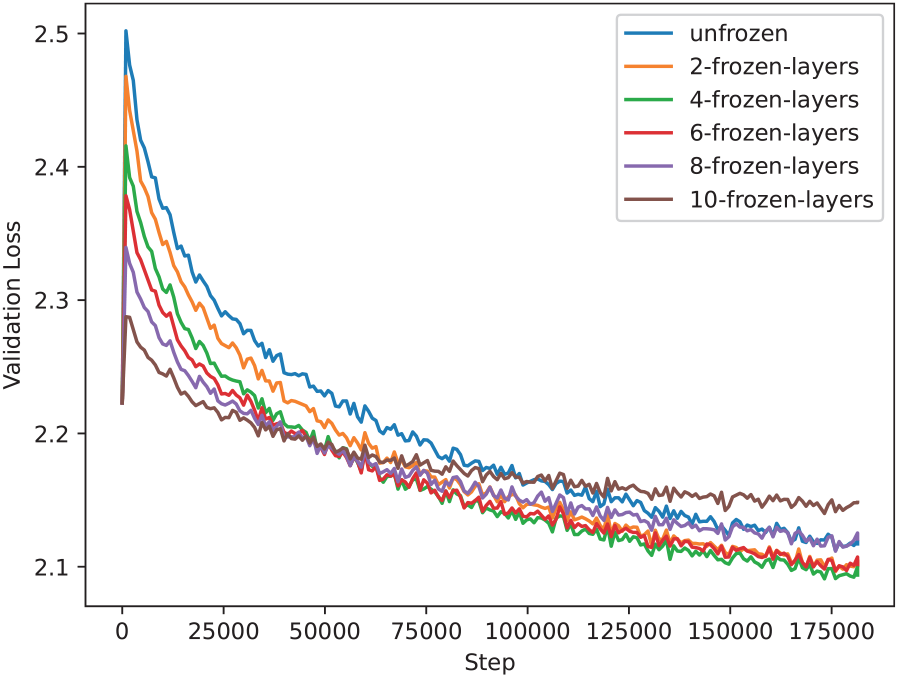
Training loss on the validation set of 24,600 OMBB protein sequences for coarse-tuning. For all models the cross-entropy loss on the OMBB validation-set sequences decreases and plateaus after 200 epochs (150000 steps). We see that the validation loss of the model with 4 frozen layers is the lowest, yet we found that the transfer learning tasks with the embeddings by these models were best with the non-frozen (0 frozen layers) model.

We quantify how well contacts are predicted on the test set of structurally characterized domains. The gold standard is the binary contact assignment calculated from the structures and depends on the Euclidean distance between their C-*α*s. The predictions are calculated from the attention matrices of the models and have continuous values between 0 and 1, as they are the logistic regression of the attention values. Figure 7 shows the ROC curves for this data. The contact predictions are far from being perfect, yet the AUC of 0.62 of the coarse-tuned model’s ROC curve is better. Also, the ROC curves have an FPR of almost 1 when their TPR is lower than 0.8, indicating that there is a large portion of the contacts that the attention matrices fail to predict.

**Fig 7.**
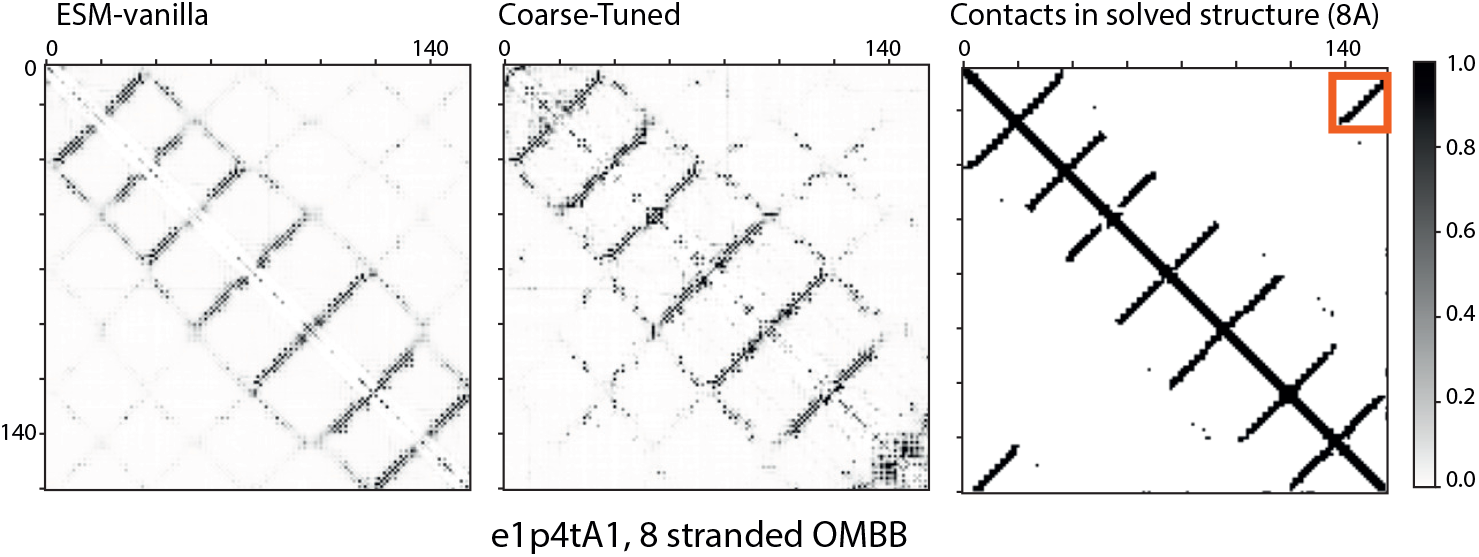
Contact map prediction and ground truth of 8Å C*α* distance for domain e1p4tA1 an 8-stranded barrel. The beta-seam contacts are highlighted by the red-box in the top right.

**Fig 8.**
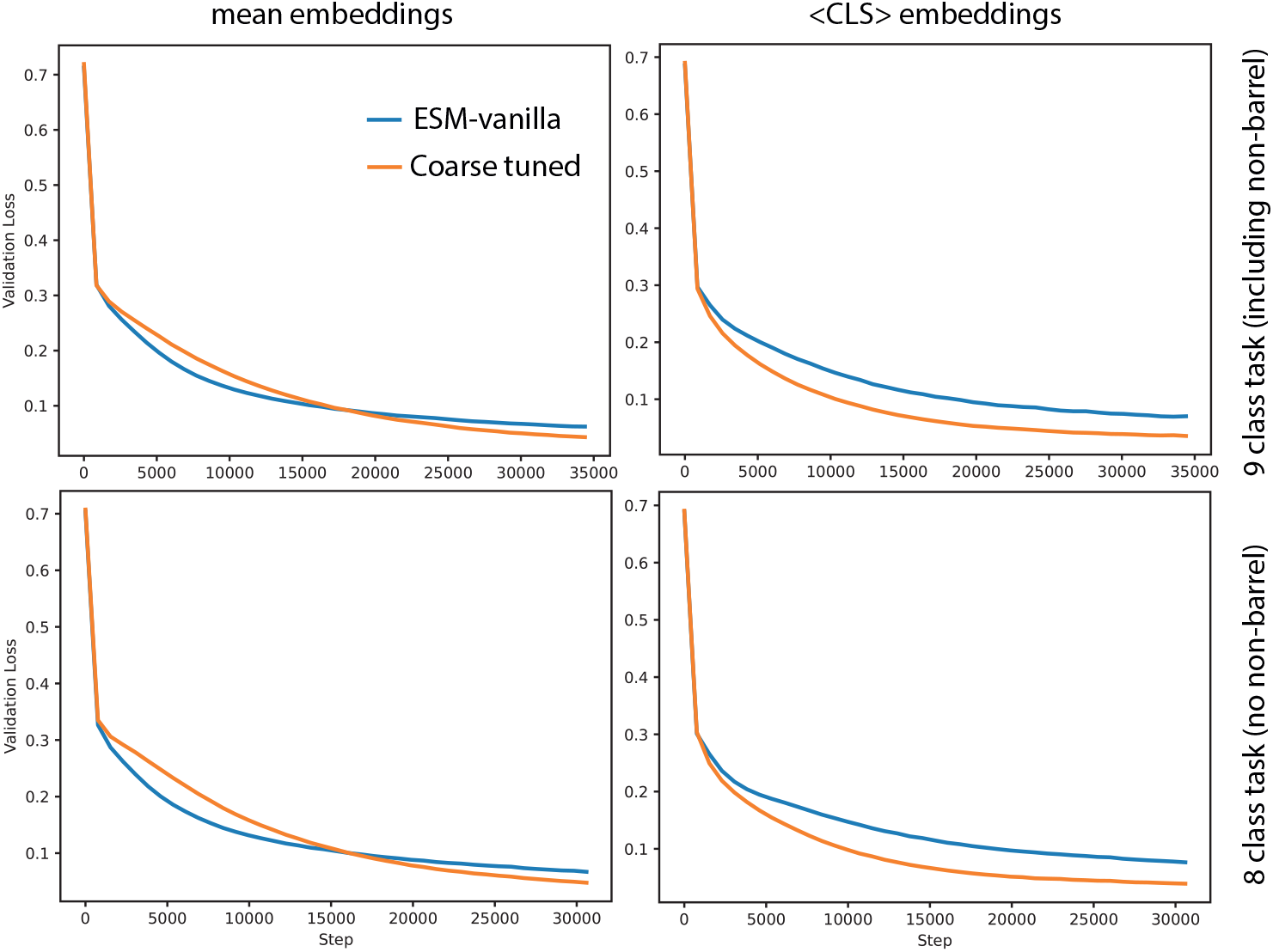
Supervised loss on validation set when training a small network to predict barrel strand number. We show loss when training on the 9 and 8 class task (with and without a non-barrel class), and when relying on the mean embeddings or the <CLS> embeddings. In all cases, the validation loss reaches a plateau.

## 3 Discussion

We propose a method that improves, or tunes, a foundation model for a class or type of proteins. We call this ‘coarse-tuning’. To coarse-tune the PLM ESM2, we used a set of ∼80K OMBB sequences and the MLM task. The resulting foundation model is better-suited for OMBBs, and we demonstrate this from multiple perspectives: the representation calculated by the coarse-tuned model on OMBB residues better captures the relationships between amino acids, transfer learning is improved, and the learned attention maps predict contacts better.

Because transfer learning can use the representations calculated by the foundation models in different ways, we tried different methods to collate the residue embeddings to a fixed-size per chain embedding and different classification models. We consider classification by dimensionality reduction and clustering using SNE with varying parameters, using *k*-NN with different distance measures (in the high-dimensional embedding). We also explore the setting in which the representation is used as input to a small network, trained in a supervised manner. In all tested cases, the classifiers using ESM-vanilla were outperformed by those using the coarse-tuned models.

To implement coarse-tuning one only needs to collect a dataset of relevant proteins and further train the foundation model using MLM. Not only is collecting a dataset far easier than labeling it, there is no reason to believe that including proteins that do not fit the inclusion criteria is a problem, as the MLM task can be learned on any protein. Thus, coarse-tuning training can be carried out on larger datasets and the model can then be used in a down-stream task, relying on transfer learning and a smaller, labeled dataset. For example, in our case we used ∼80K sequences for coarse-tuning, and only a few thousand sequences in the supervised setting. If we use the representations directly and apply *k*-NN, even a few dozens of labeled proteins suffice. Indeed, when the dataset available for supervised learning is small, transfer-learning is often considered more appropriate than fine-tuning, as the latter may lead to overfitting.

Understanding how the weights of any foundation model that embeds protein relevant data is an unmet challenge. We followed Rives et al.’s [26] analysis for ESM embeddings and visualized the relationships among the mean embeddings of different amino acids calculated for a set of proteins. When considering a set of OMBB proteins and comparing the relationships among the embeddings of different amino acids in the ESM-vanilla model and the coarse-tuned models, we found that the relationships among the weight vectors learned for each residue by a transformer model (visualized in Rives et al.) that were lost in OMBB proteins can be recovered by coarse-tuning the model on our larger set of OMBB proteins. This observation carries practical value as it suggests that if one has a set of proteins of interest, they can monitor the relationships among the embeddings of different amino acids in different models for a set of such proteins until the relationships reported for the diverse set of proteins are recovered.

That the coarse-tuned model generates embeddings that can be used to better predict the number of strands in designed sequences and the contact maps of OMBBs is a surprising feature as the sequences were designed with ESM-IF that is closely related to the ESM-vanilla foundation model. Furthermore, the coarsetuned models succeed with respect to already successful models, as ESM-vanilla based ESMFold, predicts designed OMBBs well [14].

OMBB proteins are indeed a distinct class of proteins. As they are membrane proteins, their biophysical environment of the membrane lipids is hydrophobic, in stark contrast from the water environment of soluble proteins. In fact, the outer membrane and the cytoplasmic (inner) membrane are also different, with beta-barrels generally found in the former and alpha-helical bundles in the latter [27]. OMBBs are also evolutionarily related to each other, and thought to have diverged from an ancestral beta-hairpin [8, 11, 25]. We speculate that these are the underlying reasons why coarse-tuning on a dataset of this group of proteins has led to an improved foundation model for them.

## Acknowledgments

The work was supported by BSF-NSF grant number 2022641. We thank Tejaswi Nimmagadda and Samuel Lim for help in dataset curation.

## 4 Supplementary Material

